# EIDD-3525 is a broad-spectrum antiviral against alphavirus infection that targets the nsP4 RNA-dependent RNA polymerase

**DOI:** 10.64898/2026.09.26.754626

**Authors:** Peiqi Yin, Shweta Choudhary, Sainan Wang, Bennett J. Davenport, Phong M. Truong, Zachary M. Sticher, Rebecca Krueger, Amalia Anne Cruz, McKenzie Smith, Alexander A. Kolykhalov, Dariia Vyshenska, Alexander L. Greninger, Shuli Mao, Michael G. Natchus, George R. Painter, Andres Merits, Margaret Kielian, Thomas E. Morrison

## Abstract

Mosquito-transmitted alphaviruses, including chikungunya (CHIKV), Mayaro, Eastern equine encephalitis, and Venezuelan equine encephalitis viruses, cause severe arthritic or neurologic disease in humans. There are currently no approved antiviral treatments against alphavirus infection. Here, we report that the purine ribonucleoside EIDD-3525 is a potent inhibitor of alphavirus replication *in vitro* with broad spectrum antiviral activity against the *Alphavirus* genus. Using a trans-replicase assay, we find that EIDD-3525 blocks alphavirus RNA replication. Mutational profiling suggests that EIDD-3525 targets the alphavirus nsP4 RNA-dependent RNA polymerase that is conserved across medically important alphaviruses. In a pre-clinical mouse model of CHIKV infection and disease, oral treatment with EIDD-3525 reduced viral tissue burdens and the severity of virus-induced disease. Collectively, our findings support the continued development of the purine ribonucleoside EIDD-3525 as an orally available antiviral therapeutic for use against a broad group of alphaviruses.

## INTRODUCTION

The *Alphavirus* genus in the family *Togaviridae* is a large group of enveloped, positive-strand RNA viruses [1, 2]. This genus includes numerous mosquito-transmitted human pathogens such as chikungunya virus (CHIKV), Mayaro virus (MAYV), o’nyong-nyong virus (ONNV), Ross River virus (RRV), and the Eastern (EEEV), Western (WEEV), and Venezuelan equine encephalitis (VEEV) viruses. In the last 30 years, several alphaviruses have re-emerged to cause large outbreaks. For example, in 1996-1997 ONNV caused a severe outbreak in south-central Uganda, with infection rates estimated at 45-68%, that then spread to Kenya and Tanzania [3]. Since 2004, CHIKV has spread globally and caused millions of cases of severe arthritic disease and some deaths in Africa, Asia, and the Americas [4]. Indeed, CHIKV is estimated to cause ∼35 million annual infections [5]. In 2023-2024, a novel lineage of WEEV re-emerged in Argentina after a nearly 40-year absence resulting in 217 human cases, 12 of which were fatal [6]. The re-emergence of these mosquito-transmitted alphaviruses and their capacity to cause large scale outbreaks highlights the critical need to develop antiviral therapeutics that are broadly effective against alphavirus infection.

The *Alphavirus* ∼12 kb RNA genome contains two open reading frames [2, 7, 8]. Upon delivery of the viral genome to the host cell cytosol, the first open reading frame is translated to produce nonstructural (ns) polyproteins, which are enzymatically processed by a virus-encoded ns protease to produce four ns proteins (nsP1-4) that form the viral replicase [9, 10]. The viral replicase mediates replication of a full-length negative-strand that serves as a template for replication of full-length positive-strand genomic RNAs and transcription of subgenomic mRNAs containing the second open reading frame. Translation of the subgenomic mRNA produces structural polyproteins, which are enzymatically processed by viral and host proteases. These structural proteins package the genomic RNA and assemble into new virions that ultimately bud from the host cell plasma membrane [11].

Development of antiviral strategies against alphavirus infection is an active area of investigation [12–15]. For example, promising inhibitors of viral RNA synthesis with potent *in vivo* efficacy against EEEV and VEEV infection have been developed from large scale screening and pharmacophoric efforts [16]. In addition, several groups have identified small molecule inhibitors of nsP1 capping activity and both nsP2 protease and helicase activities that are efficacious *in vitro* and *in vivo* [17–23]. The alphaviral nsP4 RNA-dependent RNA polymerase (RdRp) also is a promising antiviral target with several nucleoside analogs, such as β-D-*N*^4^-hydroxycytidine (NHC), shown to limit alphavirus replication [24, 25]. Indeed, our group found that 4′-fluorouridine (4′-FIU), a pyrimidine ribonucleoside analog, inhibits CHIKV and MAYV infection *in vitro* and *in vivo* following oral administration to mice [26].

In this study, we evaluated EIDD-3525, a purine ribonucleoside analog that has demonstrated anti-hepatitis C virus activity *in vitro* [27], for inhibitory activity against alphaviruses. We found that EIDD-3525 treatment suppressed alphavirus replication in human cells. Using a *trans*-replicase assay, we further demonstrated that EIDD-3525 treatment inhibits RNA replication and transcription by the replicase proteins from a broad group of arthritogenic and encephalitic alphaviruses. Consistent with these findings, mutational profiling suggests that EIDD-3525 targets the alphavirus nsP4 RNA-dependent RNA polymerase that is conserved across alphaviruses. Importantly, in a pre-clinical mouse model of CHIKV infection and disease, oral treatment with EIDD-3525 reduced viral tissue burdens and the severity of virus-induced disease. Taken together, our findings support the continued development of EIDD-3525 as an orally available antiviral therapeutic for use against a broad group of alphaviruses.

## RESULTS

### EIDD-3525 has broad inhibitory activity against alphavirus infection

We evaluated the antiviral potency of EIDD-3525 (**Fig 1A**) against CHIKV using CHIKV strain 181/25 engineered to express nanoluciferase downstream of the viral subgenomic promoter (CHIKV-nLuc) [26]. CHIKV 181/25 is an attenuated derivative of strain AF15561, an Asian genotype CHIKV strain isolated from an infected human in Thailand [28]. Human U-2 OS cells or normal human dermal fibroblasts (NHDF) were infected for 1 hour (h) with CHIKV-nLuc at a multiplicity of infection (MOI) of 0.1 focus forming units (FFU) per cell. The cells were then incubated with the indicated concentrations of EIDD-3525. At 24 h post-infection (hpi), cells were collected and nLuc activity was quantified. In this assay, EIDD-3525 treatment inhibited CHIKV replication with a 50% effective concentration (EC_50_) of 0.57 and 0.74 µM in U-2 OS cells and NDHF, respectively (**Fig 1B**; **Table 1**). Analysis of EIDD-3525 cytotoxicity in U-2 OS cells and NHDF showed that the half-maximal cytotoxic concentration (CC_50_) was 462 and 712 μM, respectively, resulting in high selectivity indices (SI = CC_50_/EC_50_) of ∼810-960 (**Fig 1C**; **Table 1**). We also tested the capacity of EIDD-3525 to inhibit new virus production at 24 hpi following CHIKV 181/25 infection of U-2 OS cells. Quantification of infectious virus in cell culture media by focus formation assay (FFA) revealed that EIDD-3525 inhibited virus production with an EC_50_ of 0.64 µM, similar to the EC_50_ observed in the nLuc assay (**Fig 1D**; **Table 1**). Next, we evaluated the breadth of EIDD-3525 antiviral activity against a panel of genetically distinct alphaviruses including MAYV, ONNV, RRV, Semliki Forest virus (SFV), and Sindbis virus (SINV). In all cases, treatment of U-2 OS cells 1 h after infection with EIDD-3525 inhibited virus production with EC_50_ values below that observed for CHIKV (**Fig. 1E-I**; **Table 1**), suggesting that EIDD-3525 has broad anti-alphavirus inhibitory activity.

**Figure 1.**
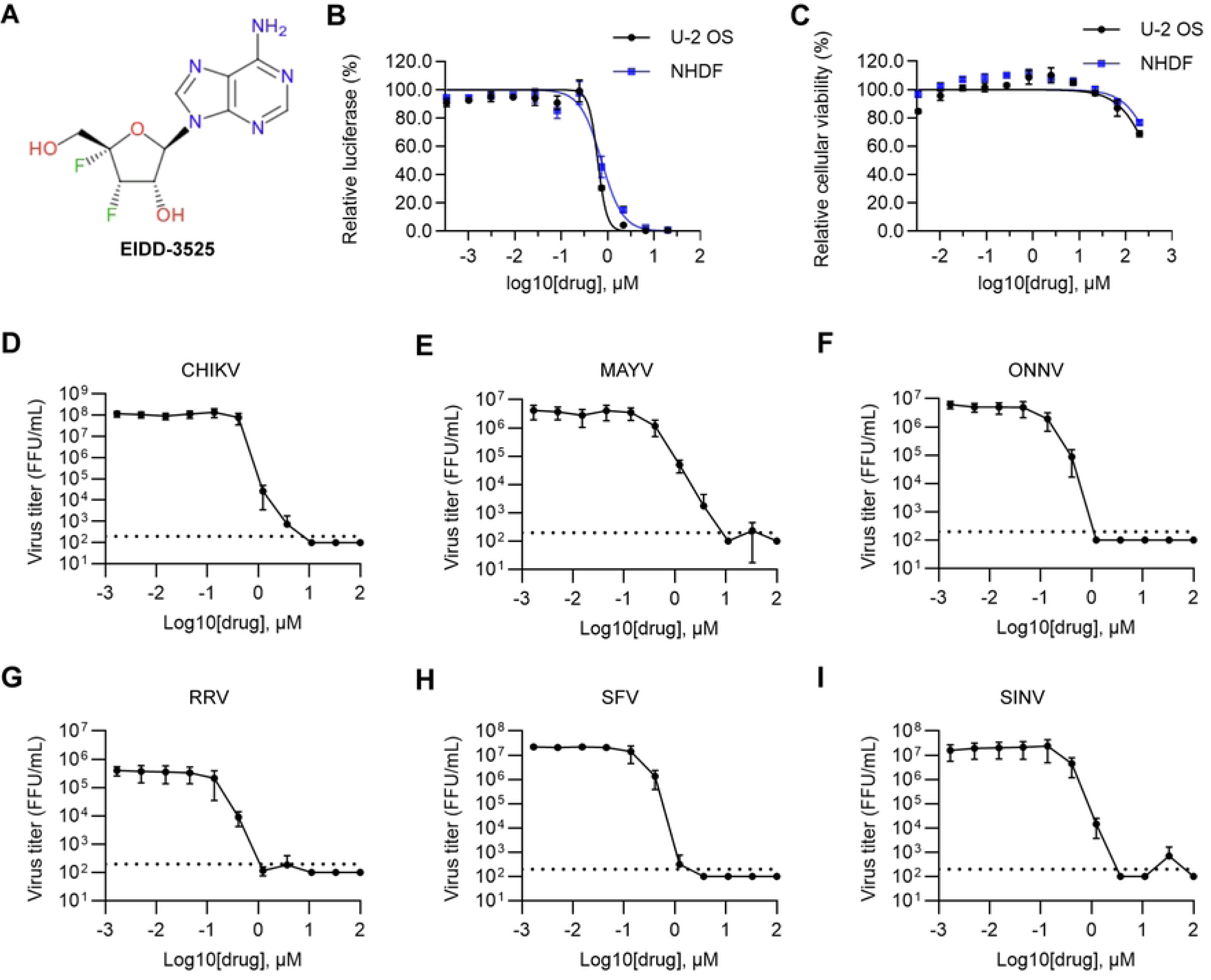
EIDD-3525 broadly inhibits alphavirus infection. **(A)** Structure of EIDD-3525. (B) U-2 OS or NHDF cells were infected with CHIKV-nLuc at an MOI = 0.1 for 1 h and treated with the indicated concentrations of EIDD-3525 or DMSO control. Nanoluciferase activity was measured at 24 hpi and normalized to that of control cells. (**C**) U-2 OS and NHDF cells were incubated with the indicated concentrations of EIDD-3525 for 24 h. Cell viability was assessed using PrestoBlue and normalized to control DMSO-treated cells. (**D-I**) U-2 OS cells were infected with CHIKV (**D**), MAYV (**E**), ONNV (**F**), RRV (**G**), SFV (**H**), or SINV (**I**) at a MOI = 0.1 for 1 h and treated with the indicated concentrations of EIDD-3525. Media were harvested at 24 hpi and titered by FFA on U-2 OS cells. The data shown represent the mean and standard deviation of three independent experiments. EC_50_ and CC_50_ values are summarized in Table 1.

**Table 1.** Summary of EIDD-3525 inhibition of alphavirus infection.

| Virus or replicon | EC<sub>50</sub> [μM] | 95% CI [μM] | Cell line | Assay readout |
| --- | --- | --- | --- | --- |
| CHIKV-nLuc | 0.57 | 0.41 to 0.79 | U-2 OS | Luciferase reporter |
| CHIKV-nLuc | 0.74 | 0.57 to 0.96 | NHDF | Luciferase reporter |
| CHIKV | 0.64 | 0.34 to 1.21 | U-2 OS | Virus titration |
| MAYV | 0.32 | 0.14 to 0.72 | U-2 OS | Virus titration |
| ONNV | 0.09 | 0.05 to 0.16 | U-2 OS | Virus titration |
| RRV | 0.12 | 0.06 to 0.27 | U-2 OS | Virus titration |
| SFV | 0.16 | 0.11 to 0.23 | U-2 OS | Virus titration |
| SINV | 0.39 | 0.15 to 1.09 | U-2 OS | Virus titration |

The CC<sub>50</sub> in U-2 OS cells was 462.24 μM (95% CI: 332.15 to 686.58 μM). The CC<sub>50</sub> in NHDF cells was 712.48 μM (95% CI: 483.16 to 1191.84 μM).

### EIDD-3525 inhibits alphavirus RNA replication

To begin to dissect the mechanism by which EIDD-3525 inhibits alphavirus replication, we added 1 µM EIDD-3525 to U-2 OS cells at times prior to and during CHIKV infection (**Fig 2A**), and quantitated the effect on virus production at 14 hpi (**Fig 2B**). Pre-treatment of U-S OS cells with EIDD-3525 for 1.5 h followed by removal of the compound prior to infection (MOI = 3 FFU/cell) did not reduce new virus production compared with cells treated with vehicle (DMSO) alone (**Fig 2B**). Similarly, simultaneous addition of CHIKV and EIDD-3525 to cells or pre-mixing EIDD-3525 with CHIKV particles and incubating for 1 h before infection of cells, followed by removal of the compound after 1 h of virus adsorption, caused no reduction in virus production (**Fig 2B**). In contrast, either addition of EIDD-3525 to cell cultures 1.5 h prior to infection (-1.5) or simultaneous addition of CHIKV and EIDD-3525 to cells (0 h), followed by subsequent maintenance of cells in the presence of EIDD-3525 reduced virus production by ∼3 logs (**Fig 2B**). In addition, inhibition of virus production was reduced ∼1 log when EIDD-3525 was added to cells at 4 hpi. No decrease in virus production was observed when EIDD-3525 was added at 8 hpi (**Fig 2B**). Collectively, these findings suggest that EIDD-3525 does not target a host pathway, alter CHIKV particles, block CHIKV attachment and entry, or limit CHIKV egress. Instead, these findings suggest that EIDD-3525 blocks viral RNA or protein synthesis.

**Figure 2.**
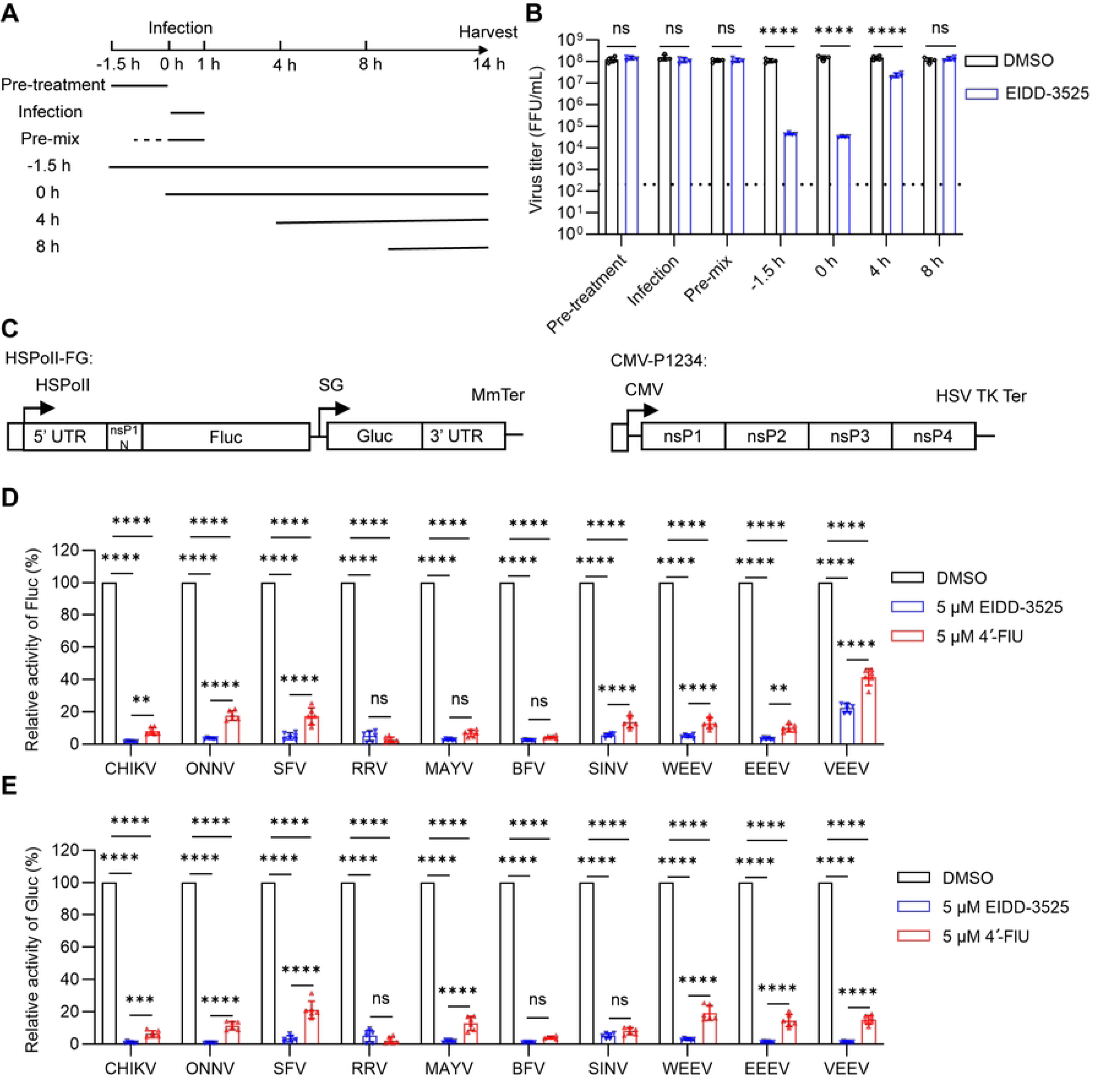
EIDD-3525 blocks alphavirus RNA replication. (**A**) Schematic of time of addition experiment. (**B**) U-2 OS cells were infected with CHIKV at MOI = 3 for 1 h. 1 µM EIDD-3525 was added as indicated. The media were harvested at 14 hpi and infectious virus was quantified by FFA. (**C**) Diagram of expression plasmids for the *trans*-replicase assay, showing template RNA (left) and replicase protein expression constructs (right). Labels indicate HSPolI, a truncated promoter (residues -211 to -1) for human RNA polymerase I; MmTer, a terminator for RNA polymerase I in mice; CMV, immediate early promoter of human cytomegalovirus; HSV TK Ter, the herpes simplex virus thymidine kinase terminator. (**D** and **E**) U-2 OS cells were transfected with alphavirus *trans*-replicase constructs for 4 h and treated with the indicated concentrations of EIDD-3525, 4′-FlU, or DMSO for 20 h. Fluc (**D**) or Gluc (**E**) activity was determined and normalized to that of DMSO-treated controls. The data shown represent the mean and standard deviation of three independent experiments. Statistical significance was calculated by two-way ANOVA with Sidak’s multiple comparisons test. **P < 0.01, ***P < 0.001, ****P < 0.0001, ns not significant.

To directly test the idea that EIDD-3525 blocks CHIKV RNA replication, we used a *trans*-replicase system in which expression of the polyprotein precursor of the viral replicase (P1234) drives replication of a reporter RNA template (**Fig 2C**) [29, 30]. In this assay, firefly luciferase (Fluc) activity represents replication of the viral RNA genome whereas Gaussian luciferase (Gluc) activity represents transcription of the viral subgenomic mRNA. Moreover, the generation of *trans*-replication assays for a variety of alphaviruses [31] allowed us to further evaluate the breadth of EIDD-3525 inhibitory activity. U-2 OS cells were transfected with 10 distinct alphavirus *trans*-replicase constructs for 4 h and treated with 5 µM EIDD-3525 or DMSO. At 20 h post-treatment, cell lysates were collected and Fluc and Gluc activity were quantified and normalized to that of DMSO-treated control cells. For comparison, cells were treated with 5 µM 4′-FlU which we previously demonstrated inhibits CHIKV RNA replication by targeting the nsP4 RNA-dependent RNA polymerase [26]. EIDD-3525 potently inhibited both CHIKV RNA replication (**Fig 2D**) and subgenomic mRNA transcription (**Fig 2E**), with activity comparable or better than that of 4′-FlU. Moreover, EIDD-3525 treatment inhibited both replication and transcription mediated by the replication complexes of all alphaviruses tested, including the encephalitic alphaviruses EEEV, VEEV, and WEEV, and this inhibition was more potent than that observed with 4′-FlU treatment (**Fig 2D-E**). Of note, EIDD-3525 inhibition of VEEV genome replication was less potent than that observed with the other alphaviruses tested.

### An amino acid substitution in the nsP4 RNA-dependent RNA polymerase influences CHIKV sensitivity to EIDD-3525

To further elucidate the mechanism of action of EIDD-3525, we passaged the attenuated CHIKV 181/25 strain in U-2 OS cells in the presence of EIDD-3525 or DMSO (**Fig 3A**). U-2 OS cells were inoculated with CHIKV 181/25 (MOI = 1 FFU/cell) and cultured in the presence of 0.5-0.75 μM EIDD-3525 (i.e., at or above the EC_50_) or DMSO for 16 h. After each passage, virus production was quantitated and the MOI was adjusted for each subsequent passage. In 6 independent passage series, virus titers were reduced by ∼100-1,000-fold at passage 2-3 in cells cultured with EIDD-3525 compared with 3 independent DMSO controls (**Fig 3B**). Inhibition of all 6 EIDD-3525 passage series was less marked by passage 4-5, with passage series #3 and #4 showing titers similar to those of DMSO treated control cells by passage 5. One additional round of passage was performed, RNA was isolated from the virus population of each passage series, reverse-transcribed, and whole genome sequencing (WGS) of CHIKV genomes was performed by Illumina deep sequencing (**Fig 3A**). A single amino acid change in nsP4 (F375Y) was detected at high allele frequency (93.3-97.4%) in EIDD-3525 passage series #2, #3, and #4 (**Fig 3C**; **Table 2**); these showed the highest virus titers at passage 5 among the 6 independent EIDD-3525 passage series (**Fig 3B**). Importantly, nsP4 F375Y was not detected in viral genomes isolated from virus passaged in the presence of DMSO or from EIDD-3525 passage series #1, #5, and #6, which remained more sensitive to EIDD-3525 inhibition (**Fig 3C**). The nsP4 F375Y change was the only mutation detected in passage series #4 virus. A few other mutations were detected in EIDD-3525 passage series #2 and #3 viruses; however, these were either synonymous changes or a change in the E2 glycoprotein (**Table 2**). Because EIDD-3525 potently inhibited viral RNA replication and transcription in the *trans*-replication assay (**Fig 2C-E**), which lacks the viral structural genes (**Fig 2C**), we considered E2 an unlikely target of EIDD-3525, although the presence of the E2 R36S mutation in passage series #2 might explain its lower growth. As shown in **Fig 3D-F**, nsP4 F375 is in the palm domain of nsP4 near the GDD active site and is conserved across the *Alphavirus* genus. 4′-FlU is a similar pyrimidine ribonucleoside that inhibits CHIKV RNA replication and subgenomic mRNA transcription by targeting nsP4 [26]. However, the F375Y amino acid substitution that arose in virus passaged in the presence of EIDD-3525 is distinct from the nsP4 amino acid changes Q192L and C483Y detected in CHIKV 181/25 that was passaged in the presence of 4′-FlU (**Fig 3D-E**) [32].

**Figure 3.**
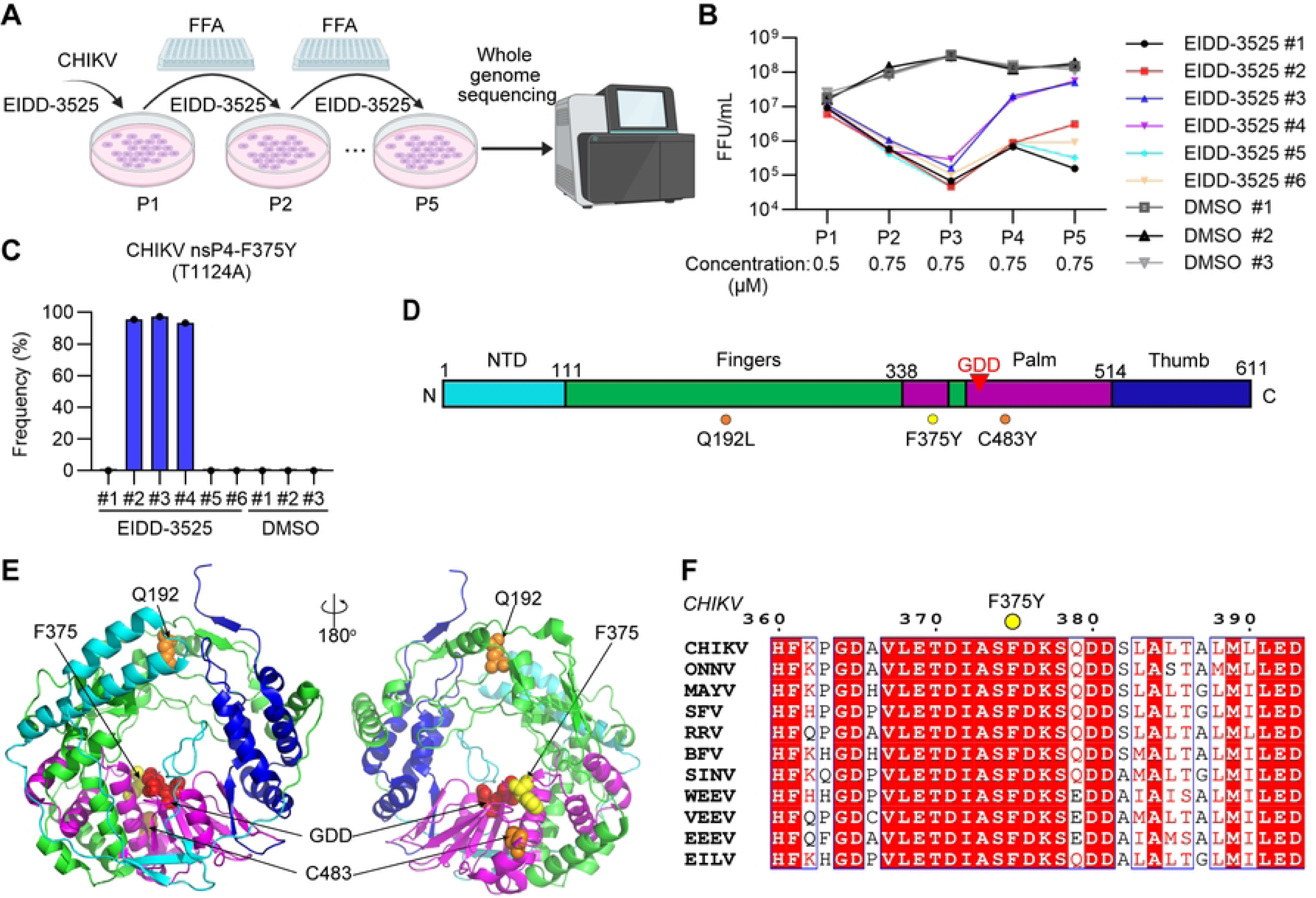
EIDD-3525 selects for amino acid changes in the CHIKV RdRp. (**A**) Schematic overview of the *in vitro* selection workflow for CHIKV under compound pressure. Figure created in BioRender. Yin, P. (2025) https://BioRender.com/ifpcoki. (**B**) Viral replication dynamics across independent CHIKV lineages subjected to serial passaging in U-2 OS cells under the indicated concentrations of EIDD-3525 or DMSO. Viral titers were quantified by FFA after each passage. (**C**) Mutation frequencies of CHIKV nsP4-F375Y (T1124A) in viral populations from six EIDD-3525 or three DMSO treated lineages after passage 6, as determined by WGS. (**D**) Schematic of the CHIKV nsP4 polymerase. Location of amino acid changes identified with EIDD-3525 (F375Y) is shown as a yellow circle. Amino acid changes previously identified against 4′-FlU are shown as orange circles. The GDD active site is denoted with red triangle. (**E**) Locations of residues that influence compound sensitivity in a 3D-model of the CHIKV nsP4 polymerase. The position of F375, the site of the amino acid change identified in virus passaged in the presence of EIDD-3525 (F375Y), is shown as yellow spheres. Sites associated with sensitivity to 4′-FlU are shown as orange spheres. The GDD active site is shown as red spheres. The NTD, fingers, palm, and thumb domains are colored cyan, green, magenta and blue, respectively. (**F**) Alignment of alphavirus nsP4 sequences around residue F375, which is indicated by a yellow dot. Swiss-Prot accession numbers: CHIKV: A4L7I2, ONNV: P13886, MAYV: Q8QZ73, SFV: P08411, RRV: P13887, BFV: P87515, SINV: P03317, WEEV: P13896, VEEV: P36328, EEEV: Q306W6. GenBank accession number of Eilat virus (EILV): BFZ80397.

**Table 2.** Summary of mutations with frequency >10% in CHIKV lineages.

| Sample | Frequency (%) | Change type | AA change <sup>a</sup> | Nucleotide change |
| --- | --- | --- | --- | --- |
| EIDD-3525 lineage #1 | 23.36 | nonsynonymous | nsP3: A137V | nsP3: C410T |
| EIDD-3525 lineage #2 | 95.41 | nonsynonymous | nsP4: F375Y | nsP4: T1124A |
|  | 95.94 | nonsynonymous | E2: R36S | E2: C106A |
| EIDD-3525 lineage #3 | 97.37 | nonsynonymous | nsP4: F375Y | nsP4: T1124A |
|  | 96.05 | synonymous | nsP4: A270A | nsP4: A810C |
|  | 91.72 | synonymous | E2: I101I | E2: C303T |
| EIDD-3525 lineage #4 | 93.35 | nonsynonymous | nsP4: F375Y | nsP4: T1124A |
| EIDD-3525 lineage #5 | 20.63 | nonsynonymous | nsP2: T794I | nsP2: C2381T |
|  | 17.33 | nonsynonymous | nsP2: G8V | nsP2: G23T |
|  | 11.11 | nonsynonymous | E2: G279E | E2: G836A |
| EIDD-3525 lineage #6 | 24.3 | nonsynonymous | nsP3: Q328L | nsP3: A983T |
|  | 21.51 | nonsynonymous | nsP4: M479L | nsP4: A1435C |
|  | 10.17 | nonsynonymous | nsP2: T5A | nsP2: A13G |
|  | 10.09 | nonsynonymous | nsP2: G8C | nsP2: G22T |

<sup>a</sup> Shown are the mutations present in the indicated independent lineages after five passages in the presence of EIDD-3525.

To determine the effect of the nsP4-F375Y substitution on EIDD-3525 sensitivity, we introduced it de novo into the attenuated CHIKV 181/25 genome and measured the effect on inhibition by EIDD-3525. In U-2 OS cells, CHIKV 181/25 harboring nsP4-F375Y displayed reduced sensitivity to EIDD-3525 compared with WT CHIKV 181/25, with a ∼2-3-fold increase in the EC_50_ (**Fig 4A**). Notably, in our CHIKV *trans*-replication assay, nsP4-F375Y had no impact on RNA replication in the absence of EIDD-3525 while an nsP4-F375A substitution abolished CHIKV RNA replication, suggesting an essential role for nsP4 position 375 in viral replication (**Fig 4B**). In the presence of EIDD-3525, nsP4-F375Y rendered CHIKV RNA replication less sensitive to inhibition, again resulting in a ∼2-3-fold increase in the EC_50_ (**Fig 4C**). Given the conservation of nsP4 F375 (**Fig 3F**), we used our panel of alphavirus *trans*-replication assays to test if an F375Y substitution in nsP4 influenced the sensitivity of other alphaviruses to EIDD-3525 inhibition. U-2 OS cells were transfected with WT or nsP4 F375Y (or F374Y) EEEV, VEEV, and WEEV replicase constructs for 4 h and the indicated concentrations of EIDD-3525 were added. Fluc and Gluc activities were measured 20 h after transfection, and normalized to those of DMSO-treated controls. EEEV, VEEV, and WEEV nsP4-F375Y/F374Y RNA replication and transcription showed decreased sensitivity to EIDD-3525 inhibition compared with their WT counterparts, with increases in the EC_50_ ranging from 3.5-23.4-fold (**Fig 4D-F**). Finally, we investigated the effect of the nsP4-F375Y substitution on CHIKV sensitivity to other antiviral ribonucleoside analogs. In U-2 OS cells, CHIKV 181/25 (WT) and CHIKV 181/25 harboring nsP4-F375Y displayed similar sensitivities to inhibition following treatment with either 4′-FlU or NHC (**Fig 5A**). Notably, CHIKV 181/25 harboring nsP4-C483Y or nsP4-Q192L, which decrease sensitivity to 4′-FlU [32], displayed increased sensitivity to inhibition by EIDD-3525 (**Fig 5C**).

**Figure 4.**
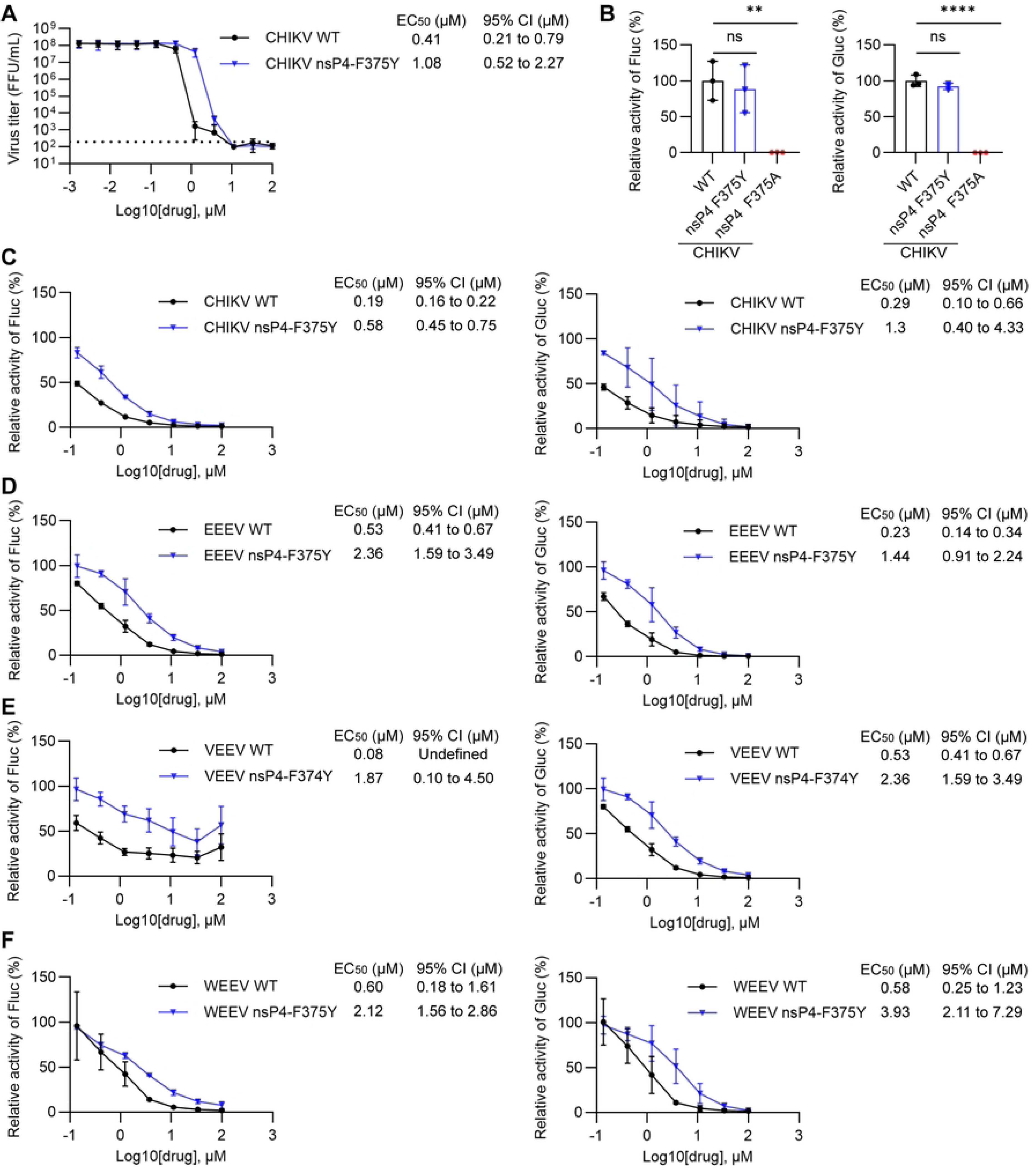
NsP4-F375Y influences sensitivity to EIDD-3525. (**A**) The substitution nsP4-F375Y was introduced into the CHIKV 181/25 genome, and the resulting virus stock was assessed for sensitivity to EIDD-3525. Viral production was measured following treatment with the indicated concentrations, as described in Fig. 1D. (**B**) U-2 OS cells were transfected with CHIKV *trans*-replicase constructs harboring the indicated nsP4 mutation for 4 h. Fluc and Gluc activities were measured 20 h after transfection and normalized to CHIKV WT. (**C-F**) U-2 OS cells were transfected with WT and mutant CHIKV (**C**), EEEV (**D**), VEEV (**E**) and WEEV (**F**) replicase constructs for 4 h and the indicated concentrations of EIDD-3525 were then added. Fluc and Gluc activities were measured 20 h after transfection and normalized to those of DMSO-treated controls. The EC_90_ values and 95% confidence interval (CI) are shown to the right. The data shown in this figure represent the mean and standard deviation of three independent experiments. **P < 0.01, ****P < 0.0001, ns not significant.

**Figure 5.**
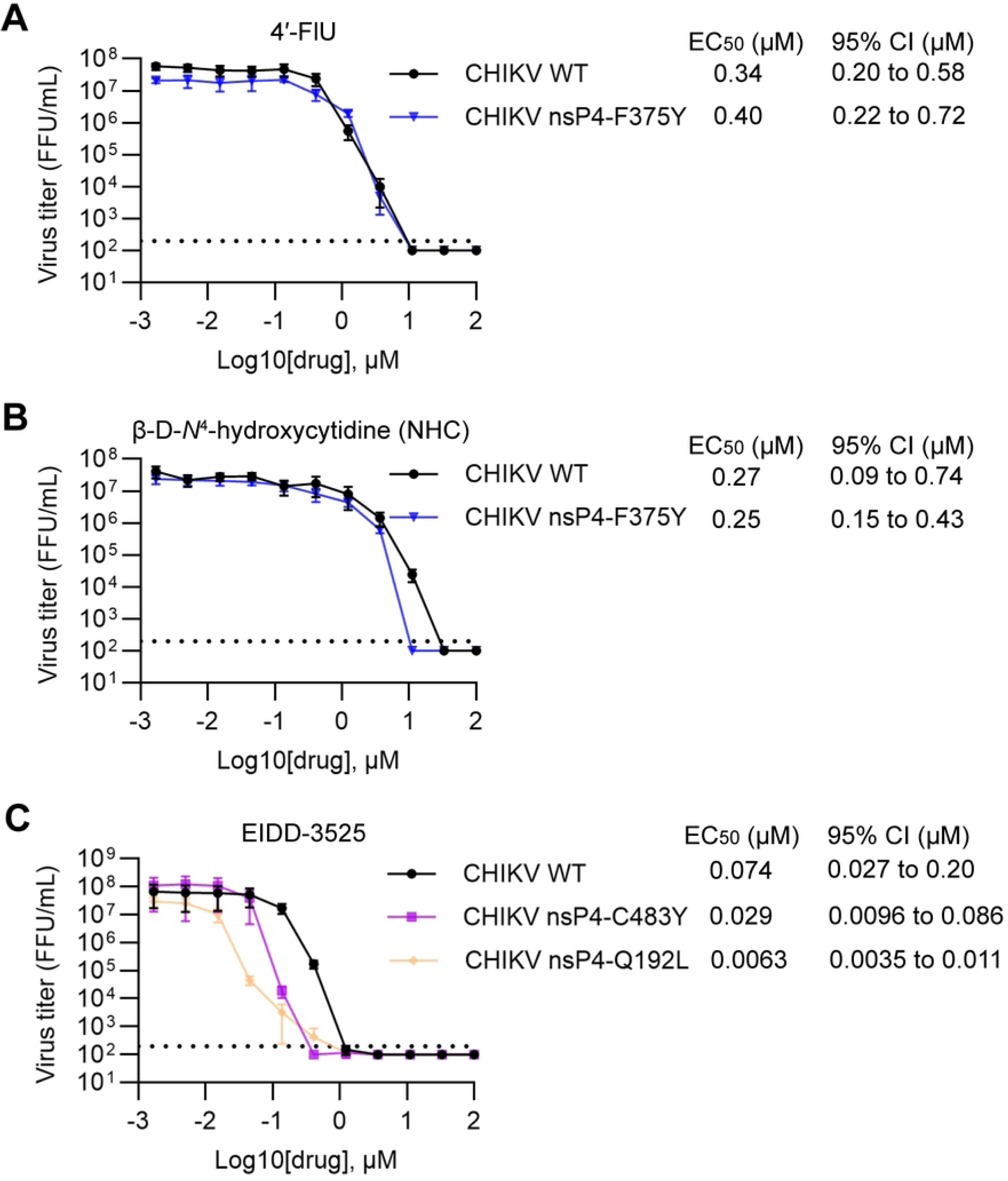
Susceptibility of CHIKV encoding nsP4 amino acid substitutions to inhibition by nucleoside analogs. (**A** and **B**) U-2 OS cells were infected with CHIKV WT or nsP4-F375Y mutant virus at a MOI = 0.1 for 1 h and treated with the indicated concentrations of 4′-FlU (A) or NHC (B). Media were harvested at 24 hpi and infectious virus was quantified by FFA. (C) U-2 OS cells were infected with CHIKV WT or viruses containing the indicated nsP4 substitutions at a MOI = 0.1 for 1 h and treated with the indicated concentrations of EIDD-3525. Media were harvested at 24 hpi and infectious virus was quantified by FFA. The EC_50_ values and 95% confidence intervals (CI) are shown to the right. The data shown represent the mean and standard deviation of three independent experiments.

### Oral treatment of mice with EIDD-3525 limits CHIKV infection and disease

To evaluate the *in vivo* tolerability and tissue distribution of EIDD-3525 and its active 5′-triphosphate metabolite, EIDD-3667, 4-week-old WT C57BL/6 mice were treated once daily (q.d.) with 300 mg/kg of EIDD-3525 or vehicle only (60% PEG 400; 40% water) by oral gavage (p.o.) (**Fig S1A**). During the one-week treatment, mice treated with EIDD-3525 gained weight similar to vehicle treated control mice (**Fig S1B**). Serum and tissues were collected on day 7 (24 h after the final dose) and the concentrations of EIDD-3525 and EIDD-3667 were determined (**Fig S1C-D**). The parent nucleoside was observed in serum 24 h after the final dose at 0.23 nml/mL. Importantly, the active EIDD-3667 form was detected in all examined tissues including musculoskeletal tissues such as ankles, wrists, and skeletal muscle that are key targets of arthritogenic alphaviruses (**Fig S1D**) [33]. However, EIDD-3667 tissue concentrations were lowest in the brain (**Fig S1D**), a key target tissue of encephalitic alphaviruses [33]. Collectively, these findings suggest that EIDD-3525 is well-tolerated *in vivo*, absorbed into blood and tissues, and anabolized to the active form at concentrations that could be efficacious against alphavirus infection.

To directly test the antiviral efficacy of EIDD-3525 against alphavirus infection *in vivo*, we used an established mouse model of CHIKV infection and disease [26]. Four-week-old wild-type (WT) C57BL/6 mice (n = 16 mice/group) were inoculated subcutaneously in the left rear footpad with 10^3^ PFU of CHIKV SL15649, an Indian Ocean lineage strain isolated from the serum of an individual infected in Sri Lanka [34]. Two hours post-inoculation, mice were treated with 300 mg/kg of EIDD-3525 or vehicle orally (p.o.) and treatment was continued once per day (q.d.). Tissues were collected for viral burden analysis at 1 (n = 8 mice/group) and 3 (n = 8 mice/group) days post-inoculation (dpi) (**Fig 6A**). At 1 dpi, after a single dose of EIDD-3525, the viral burden in the ipsilateral ankle and serum were similar to that of vehicle control mice (**Fig 6B**). At 3 dpi, the viral burden in the ipsilateral ankle and serum were reduced in mice treated with EIDD-3525 (**Fig 6B**). CHIKV infection following subcutaneous inoculation of WT C57BL/6 mice in the footpad leads to biphasic swelling of the ipsilateral foot and ankle, with an initial peak on days 2–3 post-infection and a second, more prominent peak on days 6–7 post-infection [26, 35]. Twice daily treatment of mice with 150 mg/kg EIDD-3525 (300 mg/kg/day) (**Fig 6C**), initiated at 2 h post virus inoculation, significantly reduced joint swelling on days 4-7 post-infection compared with vehicle-treated control mice (**Fig 6D**). In addition, at 7 dpi the viral burden in the ipsilateral ankle (3.4-fold; Student’s unpaired t-test), contralateral ankle (18.6-fold; Student’s unpaired t-test), ipsilateral wrist (46.8-fold); Student’s unpaired t-test), and spleen (7.4-fold; Student’s unpaired t-test) was reduced in mice treated with EIDD-3525 compared with vehicle-treated control mice (**Fig 6E**).

**Figure 6.**
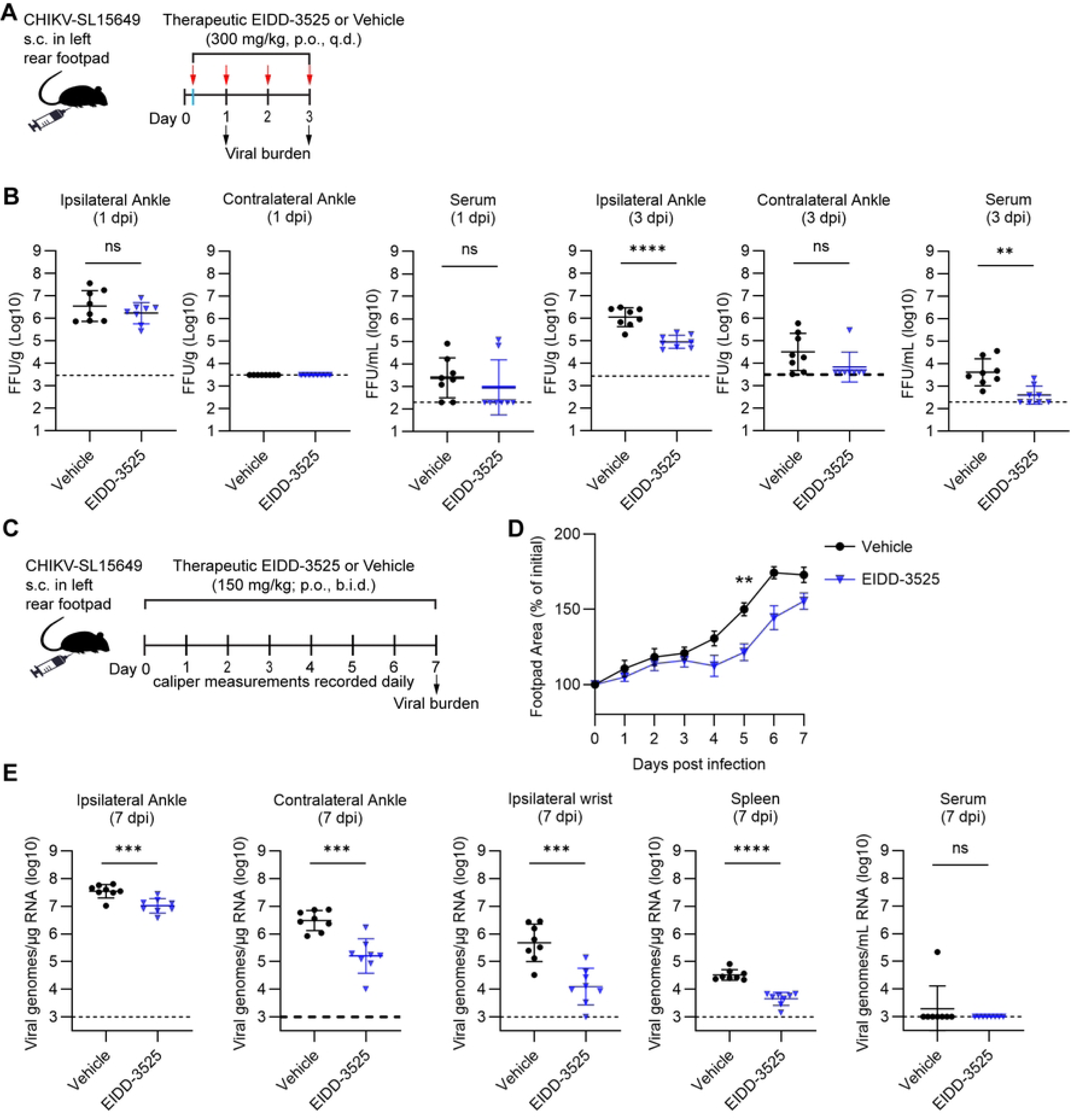
Oral treatment of mice with EIDD-3525 displays anti-viral activity against CHIKV infection. (**A**) Schematic of experimental design created with Biorender.com. WT C57BL/6 mice (n = 16 mice/group) were inoculated with 10^3^ PFU of CHIKV in the left rear footpad. At 2 h post-virus inoculation, mice were treated by oral gavage with vehicle or 300 mg/kg EIDD-3525. Treatment continued once per day until the pre-determined experimental endpoints. (**B**) At 1 (n = 8 mice/group) and 3 (n = 8 mice/group) dpi, the viral burden in the tissues indicated was quantified by FFA. Statistical significance was determined by Student’s unpaired t-test. (**C**) Schematic of experimental design created with Biorender.com. WT C57BL/6J mice (n = 8 mice/group) were inoculated with 10^3^ PFU of CHIKV in the left rear footpad. At 2 h post-virus inoculation, mice were treated by oral gavage with vehicle or 150 mg/kg EIDD-3525. Treatment continued twice per day, and the tissues were harvested at 7 days post virus challenge. (**D**) Swelling of the ipsilateral foot over time was measured by digital calipers. Statistical significance was determined by a repeated measures ANOVA with Tukey’s multiple comparison test. (**E**) At 7 dpi, the viral burden in the tissues indicated was quantified by RT-qPCR. Statistical significance was determined by Student’s unpaired t-test. \*\**P* < 0.01, \*\*\**P* < 0.001, \*\*\*\**P* < 0.0001, ns not significant.

## DISCUSSION

There is a critical need for broadly active antiviral therapies against alphaviruses, as mosquito-transmitted alphaviruses continue to emerge and re-emerge [14, 15], Here, we show that the purine ribonucleoside EIDD-3525 is a potent inhibitor of alphavirus replication. EIDD-3525 suppressed CHIKV production in human U-2 OS cells and normal human dermal fibroblasts with low cytotoxicity and inhibited RNA replication by a broad panel of arthritogenic and encephalitic alphaviruses. Mutational profiling identified the highly conserved nsP4 residue F375 as a determinant of viral sensitivity to EIDD-3525. In addition, once-daily oral treatment with EIDD-3525 reduced CHIKV tissue burdens and joint swelling in mice. These results further support the alphavirus nsP4 RNA-dependent RNA polymerase as an antiviral target [12].

Our studies indicate that EIDD-3525 acts at the viral RNA replication stage of the alphavirus replication cycle. Time-of-addition studies indicated that EIDD-3525 acts during an early post-entry step of CHIKV infection, and trans-replication assays showed that the compound inhibits both genomic RNA replication and subgenomic RNA transcription. EIDD-3525 inhibited the replication complexes of all alphaviruses tested and in this assay was generally more inhibitory than 4′-FlU at the same concentration. EIDD-3525 is metabolized to the active triphosphate EIDD-3667, however the molecular mechanism by which EIDD-3667 inhibits RNA synthesis remains to be determined. Studies with purified replication complexes will be needed to determine whether EIDD-3667 is incorporated into nascent viral RNA and whether such incorporation results in chain termination.

Our mutational profiling studies identified an nsP4 F375Y amino acid substitution in three independent CHIKV lineages that showed reduced sensitivity to EIDD-3525 treatment. The F375 residue is in the nsP4 palm domain near the conserved GDD active site and is highly conserved among alphaviruses [10, 12]. Consistent with this conservation, F to Y substitution of the corresponding residue in the nsP4 gene of EEEV, VEEV, and WEEV also decreased EIDD-3525 sensitivity in our trans-replication assay, indicating that the role of this residue in EIDD-3525 sensitivity is conserved across both arthritogenic and encephalitic alphaviruses. Replacement of F375 with alanine (A) strongly impaired CHIKV trans-replication, suggesting that this position is essential for replicase activity.

F375 is distinct from residues previously associated with decreased sensitivity to other alphavirus polymerase inhibitors. In prior studies, we found that the nsP4 substitutions Q192L and C483Y decreased CHIKV sensitivity to 4′-FlU [32]. Q192 is located in the index region of the nsP4 fingers domain, while C483 is located in the palm domain between motifs C and D [36, 37]. F375 is also distinct from the K291R substitution in the nsP4 F1 motif that is associated with CHIKV sensitivity to favipiravir, a broad-spectrum purine ribonucleoside [38]. The distinct locations of these sensitivity determinants within nsP4 suggest that EIDD-3525, 4′-FlU, and favipiravir interact differently with the alphavirus replicase or affect different steps in nucleotide recognition, incorporation, and RNA synthesis. Consistent with this idea, we found that the nsP4-F375Y substitution had no impact on CHIKV sensitivity to the antiviral ribonucleoside analogs 4′-FlU or NHC. Moreover, we also found that in contrast with the effect on sensitivity to 4′-FlU, the nsP4 substitutions Q192L and C483Y increased CHIKV sensitivity to EIDD-3525. F375Y caused only a modest decrease in EIDD-3525 sensitivity, and higher compound concentrations retained inhibitory activity against replicases harboring an nsP4 F375Y substitution. The conservation and functional importance of F375, as suggested by the profound loss of activity by a replicase harboring an nsP4 F375A substitution, may limit the amino acid residues that can be tolerated at this position. Indeed, phenylalanine and tyrosine differ by only a single hydroxyl group on the phenyl ring. The distinct sensitivity profiles of these polymerase inhibitors may also be useful in evaluating combination antiviral therapies.

Importantly, we found that oral EIDD-3525 treatment was efficacious against CHIKV infection and disease *in vivo*. Once-daily oral treatment of mice with 300 mg/kg EIDD-3525 was well tolerated over seven days, and the active triphosphate form EIDD-3667 was detected in musculoskeletal tissues that are key sites of arthritogenic alphavirus infection and pathology (30). In CHIKV-infected mice, once or twice daily treatment initiated at 2 h post-virus inoculation reduced viremia and viral burdens in joint-associated tissues and significantly decreased virus-induced foot and ankle swelling, a characteristic measure of acute musculoskeletal disease in this pre-clinical mouse model [34]. These results are consistent with our previous studies of 4′-FlU, which showed that an orally administered ribonucleoside analog targeting nsP4 can inhibit CHIKV infection and disease *in vivo* [26]. Similarly, oral treatment of mice with favipiravir or molnupiravir, another pyrimidine ribonucleoside analogue with broad spectrum antiviral activity [39], also reduced viral tissue burdens following CHIKV infection [24, 40]. In the current study, viral burdens were not detectably reduced at 1 dpi after a single dose of EIDD-3525 but were reduced at 3 dpi following continued once-daily treatment, suggesting that antiviral efficacy may depend on sufficient intracellular accumulation or maintenance of the active metabolite. Although EIDD-3525 inhibited EEEV, VEEV, and WEEV replication in trans-replication assays, we found that EIDD-3667 concentrations were lowest in the brain, an important target tissue for encephalitic alphavirus infection. Further studies will be needed to determine whether sufficient drug exposure can be achieved in the central nervous system.

In summary, our studies demonstrate that EIDD-3525 is a potent and broadly active inhibitor of alphavirus RNA replication with oral efficacy against CHIKV infection and disease in mice. Mechanistic studies identified a highly conserved residue in the nsP4 palm domain as a determinant of EIDD-3525 sensitivity across divergent alphaviruses, and this residue was distinct from sites associated with decreased sensitivity to other ribonucleoside inhibitors. Together, these findings provide new insight into alphavirus polymerase inhibition and support continued development of EIDD-3525 as a broad-spectrum antiviral therapeutic against medically important alphaviruses.

## MATERIALS AND METHODS

### Cell lines

U-2 OS cells (ATCC HTB-96) were cultured in modified McCoy’s 5A medium supplemented with 10% fetal bovine serum (FBS), 100 U/mL penicillin and 100 μg/mL streptomycin. BHK-21 cells (ATCC CCL-10) and normal human dermal fibroblasts (NHDF; ATCC PCS-201-012) were maintained in DMEM containing 10% fetal bovine serum (FBS), 100 U/mL penicillin and 100 μg/mL streptomycin. Vero cells (ATCC CCL-81) were cultured in DMEM/F-12 (Gibco 11330032) supplemented with 10% FBS, 1x nonessential amino acids (Gibco 11140– 050), sodium bicarbonate, 2 mM L-glutamine, and 100 U/mL penicillin and 100 μg/mL streptomycin. Cells were incubated at 37°C in a humidified atmosphere containing 5% CO₂. These culture conditions were described previously [26].

### Viruses

The alphaviruses and reporter viruses used in this study were generated from infectious cDNA clones as described previously [26, 41]. The viruses included CHIKV 181/25 (pSinRep5-181/25ic [42], provided by Dr. Terence S. Dermody), CHIKV-nLuc reporter virus (181/25 strain [41], MAYV (MAYV-CH-mKATE), RRV (RRV-T48-ZsGreen), ONNV (ONNV-ZsGreen), SFV (pSP6-SFV4 [43], and SINV (dsTE12Q [44], provided by Dr. Beth Levine). CHIKV strain SL15649, originally isolated from the serum collected from a patient infected with CHIKV in Sri Lanka in 2006 [34], was derived from the infectious cDNA clone and used in mouse experiments. For production of virus stocks, infectious clone plasmids were linearized and used as templates for in vitro transcription. Capped viral RNA was generated using an SP6 in vitro transcription kit (Life Technologies) and introduced into low-passage BHK-21 cells by electroporation. Culture supernatants were collected 27-30 h after electroporation, when cytopathic effects were evident, and clarified by centrifugation at 3,000 rpm for 20 min at 4 °C using a Thermo Scientific TX-1000 rotor. Virus stocks used for cell culture experiments were titrated by a focus formation assay (FFA) in U-2 OS cells, as described below.

### Site-directed mutagenesis by Gibson assembly

Mutations were introduced into the infectious cDNA clone of CHIKV strain 181/25 using Gibson assembly. PCR fragments containing the desired mutations were amplified with the primers (fragment 1-F: CCAAATCACCGACGAGTATGATGCATATCTAGACATGGTG; fragment 1-R: GAATCATCTTGGCTCTTATCATAGGAGGCTATGTCCGTTTC; fragment 2-F: GAAACGGACATAGCCTCCTATGATAAGAGCCAAGATGATTC; fragment 2-R: GCTAACGGTTTGCCCAATTTAAATAACCTTTTTAGCGGG). The 181/25 infectious clone plasmid were linearized by digestion with BsaB1 and SwaI, and the corresponding PCR products were assembled into the linearized plasmid backbones using the Gibson Assembly Master Mix (New England Biolabs, E2611) according to the manufacturer’s instructions.

### In vitro dose-response antiviral assays

The antiviral activity of EIDD-3525, 4′-FlU, and β-D-*N*^4^-hydroxycytidine (NHC; MedChemExpress, #3258-02-4), was evaluated using reporter-based and virus-yield dose-response assays adapted from those previously described [26, 32, 41]. For assays using CHIKV-nLuc, U-2 OS cells were seeded in 96-well plates at 1.5 x 10^4^ cells per well and incubated for 24 h. Cells were infected with the indicated alphavirus at a multiplicity of infection (MOI) of 0.1 in Med-A (minimum essential medium plus 0.2% BSA and 10 mM Hepes) for 1 h and washed three times with U-2 OS medium. 3-fold serial dilutions of EIDD-3525 were then added in U-2 OS medium. DMSO-treated infected cells were included as controls. At 24 h post-infection, cells were lysed, and nanoluciferase activity was measured using Nano-Glo Dual-Luciferase Reporter Assay reagents (Promega) and a Victor X5 multilabel plate reader (PerkinElmer). For virus-yield assays, U-2 OS cells were seeded in 24-well plates at 7.5 x 10^4^ cells per well and incubated for 24 h. Cells were inoculated with the indicated alphavirus at MOI = 0.1 in Med-A and treated with threefold serial dilutions of EIDD-3525, 4′-FlU, or NHC under the conditions described above. At 24 h post-inoculation the culture supernatants were collected and clarified by centrifugation, and infectious virus production was quantified by FFA. Virus production and luciferase signals were normalized to those of DMSO-treated controls. Dose-response curves were fitted by nonlinear regression using a variable-slope model in Prism 10 (GraphPad). Half-maximal effective concentrations (EC_50_) and the corresponding 95% confidence intervals were calculated from the fitted curves.

### Cytotoxicity assay

The effect of EIDD-3525 on U-2 OS or NHDF cell viability was assessed using the PrestoBlue assay. Cells were seeded into 96-well plates at 1.5 x 10^4^ cells per well and cultured for 24 h. The cells were then incubated for 24 h at 37 °C in culture medium containing threefold serial dilutions of EIDD-3525 to a maximum concentration of 200 µM. Following compound treatment, PrestoBlue Cell Viability Reagent (Thermo Fisher Scientific) was added to the cells, and plates were incubated for 1 h at 37 °C. Fluorescence was measured using a Victor X5 multilabel plate reader. Cell viability was normalized to that of DMSO-treated controls, and the half-maximal cytotoxic concentration (CC_50_), together with its 95% confidence interval, was determined by variable-slope nonlinear regression in Prism 10.

### Time-of-addition studies

U-2 OS cells were seeded in 24-well plates at 7.5 x10^4^ cells per well and cultured for 24 h. Cells were inoculated with CHIKV at MOI=3 in Med-A for 1 h. The virus was removed and the cells were washed three times with complete medium and cultured at 37 °C. EIDD-3525 was added at a final concentration of 1 µM at the indicated times before, during, or after inoculation. Vehicle controls received an equivalent amount of DMSO. At 14 h post-inoculation, culture supernatants were collected and clarified by centrifugation. Infectious virus production was measured by FFA on U-2 OS cells.

### Trans-replication assays

The alphavirus trans-replication system was described previously [26, 32, 41]. The system consisted of separate plasmids encoding P1234 and a replication-competent reporter RNA template for the indicated alphavirus. The reporter templates expressed Firefly luciferase to detect viral RNA synthesis and *Gaussia* luciferase to measure the viral RNA synthesized from the SG promoter. U-2 OS cells were seeded in 48-well plates at approximately 3 x 10^4^ cells per well. After 24 h, cells were co-transfected with 250 ng of the reporter RNA template plasmid and 250 ng of the corresponding P1234 expression plasmid. Transfections were performed using Lipofectamine LTX with PLUS Reagent (Thermo Fisher Scientific) according to the manufacturer’s protocol. At 4 h post-transfection, the transfection medium was replaced with fresh medium containing 5 μΜ EIDD-3525, 5 μΜ 4′-FlU, or serial dilutions of EIDD-3525. DMSO was included as a vehicle control. Cells were incubated for an additional 20 h at 37 °C and then lysed. Firefly and *Gaussia* luciferase activities were measured using the Dual-Luciferase Reporter Assay System (Promega). Reporter activity was normalized to the vehicle-treated control. Dose-response curves were analyzed by log(inhibitor) vs response (three parameters) in Prism 10 (GraphPad), and EC_50_ values were calculated from the fitted curves.

### Sensitivity testing

U-2 OS cells were seeded in six-well plates at 2 x 10^5^ cells per well and cultured for approximately 24 h. Cells were then inoculated with the primary CHIKV stock at MOI=1 FFU/cell. At 2 h post-infection, medium containing the indicated compound or an equivalent volume of DMSO was added to independent wells. At 16 h post-infection, culture supernatants were collected, clarified by centrifugation, stored frozen, and subsequently titrated by FFA. The compound concentrations used during each passage are indicated in Fig. 3B. Viruses collected after passage 5 (P5) were used for viral RNA extraction. Viral RNA was reverse transcribed, and the resulting material was subjected to whole-genome sequencing as described below.

### Whole-genome sequencing and data analysis

Viral whole-genome sequencing of CHIKV samples was performed using a metagenomic next-generation sequencing approach, as previously described [45]. Libraries were sequenced on NextSeq 2000 with 2 x 150 bp or 1 x 100 bpread format. Raw reads were trimmed and quality filtered with fastp (v0.23.4) [46] (--cut_mean_quality 20 --cut_front --cut_tail --length_required 20 --low_complexity_filter --trim_poly_g --trim_poly_x). Filtered reads were then used for variant calling using the RAVA workflow (default parameters) and the CHIKV 181/25 strain (MK028839.1) as a reference (https://github.com/greninger-lab/RAVA_Pipeline/tree/2025-08-14_AECM_MK-PY_C3-4_CHIKV_publication) [47].

### Biosafety

All experiments were reviewed and approved by the Institutional Biosafety Committee of the University of Colorado Anschutz under protocol 1051. All work involving infectious CHIKV SL15649 was conducted in approved Biosafety Level 3 (BSL-3) and animal BSL-3 laboratories.

### Mouse experiments

WT C57BL/6J mice were obtained directly from The Jackson Laboratory. All experiments were performed in 4-week-old male and female mice. Eight mice per group (four males and four females) were used for all studies. Mice were group housed in a facility with a 14 h light and 10 h dark cycle, ambient temperature of 72 ± 2 °F, and ambient humidity of 40 ± 10%. No statistical methods were used to predetermine mouse sample sizes, but our sample sizes are similar to those reported in our prior antiviral efficacy studies [26]. Mice were randomly assigned to experimental groups, anesthetized with isoflurane vapors, and inoculated in the left rear footpad with a 10 μL volume containing 10^3^ plaque-forming units (PFU) of CHIKV SL15649 diluted in phosphate buffered saline (PBS) with 1% fetal bovine serum (FBS) using a Hamilton syringe and 30 G needle. Mice were treated therapeutically with 150 mg/kg EIDD-3525, 300 mg/kg EIDD-3525 or vehicle (60% PEG 400; 40% water) by oral gavage (p.o.) once (300 mg/kg) (q.d.) or twice (150 mg/kg) (b.i.d.) daily starting 2 h post-virus inoculation. At 1, 3, and 7 dpi, sera were collected, mice were intracardially perfused with 10 mL of 1x PBS, and tissues were collected in 1x PBS/1% FBS/1x Ca^2+^Mg^2+^ for viral burden analysis by FFA. Tissues were homogenized using MP Biomedicals FastPrep-24 Classic (1 cycle of 30 seconds at 4.0 m/s). Alternatively, tissues were collected in TRIzol Reagent (Invitrogen), homogenized using MP Biomedicals FastPrep-24 Classic (1 cycle of 20 seconds at 5.5 m/s), and viral burden analysis was performed by RT-qPCR as outlined below. Swelling measurements were assessed daily using digital calipers as CHIKV SL15649 infection results in bi-phasic swelling of the ipsilateral foot and ankle in C57BL/6 mice, which can be used as an indicator of acute CHIKV joint pathology in the mouse model [48].

The tissue distribution profiling of EIDD-3525 and it’s active 5′-triphosphate metabolite, EIDD-3667, in mice was performed by treating mice with 300 mg/kg EIDD-3525, p.o., q.d. for seven days. Tissue samples were collected 24 h after the final dose, weighed, and immediately snap frozen in liquid nitrogen [26]. Aliquots of serum were mixed with 70% acetonitrile that included EIDD-3752 (^13^C5-labeled EIDD-3525) as the internal standard. Samples were vortexed for one min and centrifuged at 15,000 rpm for 5 min. The resulting supernatants were transferred to HPLC vials and HPLC separation was performed on an Agilent 1260 system (Agilent Technologies, Santa Clara, CA, USA) equipped with an autosampler, column oven, UV lamp, and binary pump. An Eclipse XDB-CN (100 x 4.6 mm, 3.5 µm) column (Thermo Fisher Scientific, Waltham, MA, USA) was used for the separation of EIDD-3525. Mobile phase A consisted of 25 mM ammonium formate buffer in HPLC grade water and mobile phase B consisted of acetonitrile. A 4-min gradient HPLC method was used. Mass spectrometry analysis was performed on a Triple Quad 7500 Mass Spectrometer (Sciex, Framingham, MA, USA) using positive mode electrospray ionization (ESI) in multiple reaction monitoring (MRM) mode. Data analysis was performed using SciexOS Software (Sciex, Framingham, MA, USA). PK parameters were calculated using Phoenix WinNonLin 8.5 (Build 8.5.1.3; Certara, Princeton, NJ) using the non-compartmental analysis tool. Samples of frozen animal tissue were extracted with cold (4 °C) (1:1) methanol: 50 mM EDTA in water that included EIDD-3752, EIDD-3761, and ^13^C10 ^15^N5 ATP as internal standards by homogenization in a BeadRuptor Elite bead mill outfitted with a cryo cooling unit (Omni, Kennesaw, GA, USA). To remove large solids, the homogenate was centrifuged for 5 min at 10,000 rpm. The supernatant was transferred to a micro-centrifuge tube, and an equal sample volume of methanol was added to each tube and vortexed. Samples were then centrifuged 5 min at 15,000 rpm to remove any remaining solids. The remaining supernatant was transferred to HPLC vials containing an equal sample volume of HPLC grade water and analyzed via LCMS MS. During analysis, samples were maintained at 4 °C. A SeQuant ZIC-pHILIC (100 x 4.6 mm, 5 μm) column (Merck Millipore, Burlington, MA, USA) was used for the separation of EIDD-3525, EIDD-3667, and ATP. Mobile phase A consisted of 50 mM ammonium bicarbonate buffer in HPLC grade water, pH-adjusted to 9.8, and mobile phase B consisted of acetonitrile. An 11.5-min isocratic HPLC method at 29% mobile phase A was performed to separate the analytes. In cases where there was an interference for EIDD-3525, an Eclipse XDB-CN (100 x 4.6 mm, 3.5 μm) column (Agilent Technologies, Santa Clara, CA, USA) was used for the analysis of EIDD-3525. For this method, mobile phase A consisted of 25 mM ammonium formate buffer in HPLC grade water and mobile phase B consisted of acetonitrile. A 4-min gradient HPLC method was used, starting with a 2-min hold at 15% B, followed by a 0.5-min gradient to 50%B, and then a return to starting conditions for 1.5 min. Mass spectrometry was performed on a Triple Quad 7500 or QTrap 7500 mass spectrometer (Sciex, Framingham, MA, USA) using negative and positive mode electrospray ionization (ESI) in multiple reaction monitoring (MRM) mode. Data analysis was performed using SciexOS Software (Sciex, Framingham, MA, USA).

### Focus formation assay (FFA)

To measure the virus titers from *in vitro* assays, U-2 OS cells were seeded in 96-well plates at 1.5 x 10^4^ per well and cultured for 24 h. Cells were inoculated with 10-fold serial dilutions of virus samples in Med-A (minimum essential medium supplemented with 0.2% BSA and 10 mM HEPES) for 2 h, and then medium was replaced with an overlay containing 1% carboxymethylcellulose, 2% FBS, and 10 mM HEPES in modified Eagle’s medium and cells cultured for 18 h. Cells were fixed with 1% PFA for 1 h, washed with PBS, permeabilized and immunostained in Perm wash (PBS containing 0.1% saponin and 0.1% BSA). CHIKV, SFV, ONNV, and MAYV-infected cells were detected using the E2-specific monoclonal antibody E2-1 [43]. RRV-infected cells were detected using a rabbit polyclonal antibody against the SFV capsid [49], and SINV-infected cells were detected using the R2 and R6 monoclonal antibodies against SINV E1 and E2 [50]. After overnight incubation and repeated washes with PBS, cells were treated with secondary goat anti-mouse or anti-rabbit IgG horseradish peroxidase-conjugated antibody diluted 1:2000 in Perm Wash. Foci were developed using TrueBlue Peroxidase substrate (#5510-0030, Seracare) and quantified on an ImmunoSpot S6 Macroanalyzer (Cellular Technologies).

Infectious virus in murine tissues was quantified as previously described [26], Briefly, 10-fold serial dilutions of homogenized tissues were generated and absorbed onto Vero cells in a 96-well plate for 2 h. 1% methylcellulose in minimal essential media (MEM) alpha/2% FBS/10 mM HEPES was used to overlay cells and then incubated at 37°C for 18 h. Cells were fixed with 1% PFA and then probed with CHK-11 monoclonal antibody [51] diluted in 1x PBS/0.1% saponin/0.1% BSA (Perm Wash) at 500 ng/mL. After incubation for 1 h and repeated washes with PBS, cells were treated with a secondary goat anti-mouse IgG horseradish peroxidase-conjugated antibody diluted 1:2000 in Perm Wash and foci developed and quantitated as above.

### Real-time quantitative polymerase chain reaction

Viral RNA in tissues was quantified as previously described [26]. Briefly, total RNA was extracted from tissues homogenized in TRIzol using a PureLink RNA Mini Kit (Invitrogen). Complementary DNA (cDNA) was generated from tissue-derived total RNA using random hexamer primers and SuperScript IV reverse transcriptase (Invitrogen). CHIKV copies were quantified using CHIKV-specific forward primer (5′-TTTGCGTGCCACTCTGG-3′) and reverse primer (5′-CGGGTCACCACAAAGTACAA-3′) with an internal TaqMan probe (5′-ACTTGCTTTGATCGCCTTGGTGAGA-3′), all within the nsP2 region of the genome [26]. The total number of CHIKV RNA copies was determined from a standard curve generated from samples containing 10^8^ to 10^0^ copies of *in vitro* synthesized CHIKV RNA spiked into total cellular RNA, and cDNA was synthesized as above. Samples were run and analyzed on a QuantStudio 7 Real-Time PCR system (Applied Biosystems).

### Statistical analysis

Statistical analyses were conducted using Prism 11 Version 11.0.0 (GraphPad). Depending on the experimental design, comparisons were performed using an unpaired two-tailed Student’s t test, or two-way ANOVA with multiple comparisons. The number of independent experiments or biological replicates and the statistical tests used are specified in the corresponding figure legends.

## ACKNOWLEDGEMENTS

We thank Elizabeth Sobolik of the University of Washington for sequence confirmation of the CHIKV SL15649 virus stocks used in the mouse studies. We thank the Einstein Analytical Imaging Facility for use of their instruments, and the following facility staff for expert training and technical assistance: Vera DesMarais and Andrea Briceno. Facilities at Einstein were supported in part by the Cancer Center Core Support Grant NIH/NCI P30-CA013330. We thank Dr. Jonathan Lai at Einstein for the use of his ImmunoSpot S6 Macroanalyzer.

## DATA AVAILABILITY

Data are available under BioProject accession PRJNA1306170.

## ETHICS APPROVAL

This study was performed in accordance with the recommendations in the Guide for the Care and Use of Laboratory Animals of the National Institutes of Health. Animal studies were conducted following approved institutional animal care and use committee (IACUC) protocols (#00215) of the University of Colorado School of Medicine (Assurance Number D16-00171) which has an AAALAC accredited animal care and use program (#00235). Experimental animals were humanely euthanized at defined endpoints by exposure to isoflurane vapors followed by bilateral thoracotomy.

## FUNDING STATEMENT

This work was supported by U19 grant AI171403 Project 2 to M.K., T.E.M., A.M., and to M.N., G.R.P., and A.L.G.

## CONFLICTS OF INTEREST

The funders had no role in study design, data collection and interpretation, or the decision to submit the work for publication. The content of this paper is solely the responsibility of the authors and does not necessarily represent the official views of the NIH or NIAID. G.R.P., M.G.N, S.M., and P.M.T. are coinventors on patent WO 2025/235664 which covers the composition of matter and the use of EIDD-3525 and its analogs as an antiviral treatment. This could affect their personal financial status. The other authors declare no conflicts of interest.

---

**Fig. S1. Tissue distribution of EIDD-3525.(A)** Schematic of experimental design created with Biorender.com. WT C57BL/6 mice (n = 4 vehicle; n = 8 EIDD-3525) were treated by oral gavage with vehicle only or 300 mg/kg EIDD-3525. Treatment continued once per day until the pre-determined experimental endpoint at 7 days post-treatment initiation. To assess toxicity of the treatment, mouse body weights were measured daily. At 24 h after the final treatment, tissues were collected, weighed, and analyzed. (**B**) The percentage of body weight relative to initial body weight over time. Statistical significance was determined by repeated measures ANOVA with Tukey’s multiple comparison test. ns not significant. (**C and D**) The amount of EIDD-3525 (**C**) and its active 5′-triphosphate metabolite, EIDD-3667 (**D**) in tissues was quantitated. Data show the mean ± SD of measurements on 8 mice.

